# Vascular smooth muscle-derived TRPV1-expressing progenitors are a new source of cold-induced thermogenic adipocytes

**DOI:** 10.1101/2020.06.26.174094

**Authors:** Farnaz Shamsi, Matthew D. Lynes, Mary Piper, Li-Lun Ho, Tian Lian Huang, Yu-Hua Tseng

## Abstract

Brown adipose tissue (BAT) functions in energy expenditure in part due its role in thermoregulation. The prominent capacity of BAT to enhance fuel utilization and energy expenditure makes it an attractive target for treating obesity and metabolic disorders. Prolonged cold exposure induces *de novo* recruitment of brown adipocytes and activates their thermogenic activity. However, the exact source of cold-induced brown adipocytes is not completely understood. In this study, we sought to investigate the cellular origin of cold-induced brown adipocytes using single-cell RNA sequencing. We identified two distinct types of adipocyte progenitors that contribute to *de novo* recruitment of brown adipocytes in response to cold challenge. One is the previously known Pdgfra-expressing mesenchymal progenitors and the other is a vascular smooth muscle-derived adipocyte progenitor (VSM-APC) population, which expresses the temperature-sensitive ion channel transient receptor potential cation channel subfamily V member 1 (Trpv1). Using flow cytometry and lineage tracing, we demonstrated that the Trpv1^pos^ VSM-APCs were indeed distinct from the Pdgfra^pos^ progenitors and could contribute to brown adipocytes with greater thermogenic potential. Together, these findings illustrate a landscape of thermogenic adipose niche at the single cell resolution and identify a new cellular origin for the development of brown adipocytes.

---

Brown adipose tissue (BAT) and related beige fat are specialized for energy expenditure, and thus the classical brown and inducible beige adipocytes are collectively called thermogenic adipocytes. Considering the immense capacity of BAT for energy expenditure and its role in fatty acid and glucose metabolism, strategies leading to increased mass or enhanced activity of BAT can potentially be utilized to combat obesity and its sequelae^1^. The main cell type in adipose tissue that stores or consumes energy is the mature adipocyte, and because mature adipocytes are postmitotic tissue expansion requires proliferation and differentiation of adipocyte precursor cells to form new mature adipocytes both during development and throughout life^2^. In addition to mature adipocytes, adipose tissue is composed of adipocyte progenitors, vascular cells, immune cells, neurons and other cell types which form the stromal vascular fraction (SVF) of adipose tissue. Although adipocytes play the major role in maintaining energy balance, cells in the SVF form the adipocyte niche and regulate adipose tissue function. The orchestrated function of cells in the adipose niche is essential for maintaining tissue homeostasis and metabolic health^3,4^.

During development in mice, mature adipocytes in classical BAT arise almost exclusively from precursor cells in the dermomyotome that are defined by expression of the transcription factor Pax3/7 and myogenic factor 5 (Myf5)^5-7^. This is in contrast to different white adipose tissue depots, where the majority of mature adipocytes arise from cells that lack expression of Myf5^8^. During adulthood, adipose depots undergo dynamic remodeling in response to environmental stimuli such as nutritional load and temperature^9,10^. In rodents, prolonged cold exposure increases BAT activity by enhancing the thermogenic capacity of pre-existing brown adipocytes and inducing *de novo* recruitment of brown adipocytes from the adipose niche in the SVF^2,11,12^. Previous studies has shown that progenitors expressing platelet derived growth factor receptor alpha (Pdgfra) and stem cell antigen 1 (Sca1) (Pdgfra^pos^ Sca1^pos^ APCs) can give rise to mature brown and beige adipocytes^13,14^. Lee et al. showed that cold triggers the proliferation of Pdgfra^pos^ Sca1^pos^ APCs followed by the induction of brown adipogenesis^13^. Additionally, it has been demonstrated that APCs expressing smooth muscle lineage markers, such as SMA and Myh11, can differentiate into beige, but not brown adipocytes^15,16^.

The present study reveals a novel cellular origin of thermogenic brown adipocytes which are distinct from the previously identified Pdgfra^pos^ Sca1^pos^ APCs. Using unbiased single-cell RNA-sequencing (scRNA-seq) analyses, we unravel a high degree of heterogeneity of vascular smooth muscle (VSM) cells and demonstrate a critical contribution of VSM to cold-induced brown adipogenesis. Importantly, we identify the expression of transient receptor potential cation channel subfamily V member 1 (Trpv1) as a specific characteristic of VSM-derived APCs. Trpv1 is a heat-activated ion channel predominantly expressed in nociceptor and thermosensory neurons in the dermal and epidermal layers of the skin, the oral and nasal mucosa, joints, sensory ganglia, and few discrete brain regions^17,18^. Lineage tracing experiments reveal the contribution of Trpv1^pos^ VSM-APCs to brown adipogenesis and establish these cells as a previously unknown source of *de novo* brown adipocyte recruitment in response to cold. Brown adipocytes originated from Trpv1^pos^ APCs express higher levels of thermogenic genes compared with those derived from Trpv1^neg^ lineage. These results suggest a new cell source for thermogenic adipocytes that could be used to design therapeutic strategies for obesity and its related metabolic disorders.

## scRNA-sequencing of BAT provides a high-resolution map of different cell types in brown adipose niche

To characterize the cellular composition of the BAT niche during cold-induced remodeling, we applied scRNA-seq to cells isolated from the stromal vascular fraction of BAT (BAT-SVF) from mice housed at either thermoneutral (TN: 30 °C for 1 week), room temperature (RT: 22 °C) or cold (5 °C for 2 days or 7 days). We sequenced 105,228 high-quality cells including both hematopoietic lineage positive and negative cells. Using a graph-based approach (Seurat)^19^, we assigned the cells into different clusters based on their gene expression similarities, then used known cell identity markers to classify specific cell types. After removal of all hematopoietic lineage positive cells followed by re-clustering of 24,498 non-hematopoietic cells and visualizing of the clusters using Uniform Manifold Approximation and Projection (UMAP) method^20^, we identified eight major non-immune cell types present in BAT-SVF (Figure 1a and Supplementary Table 1). This unsupervised clustering of gene expression profiles also revealed the heterogeneity within each cell type, illustrated by the presence of multiple distinct clusters for each cell type. Using a combination of known markers, we identified four sub-populations of Pdgfra-expressing adipose progenitors, vascular and lymphatic endothelial cells, vascular smooth muscle cells, pericytes, differentiating adipocytes, as well as myelinating- and non-myelinating Schwann cells (Figure 1b and Extended Data Figure 1).

**Figure 1.**
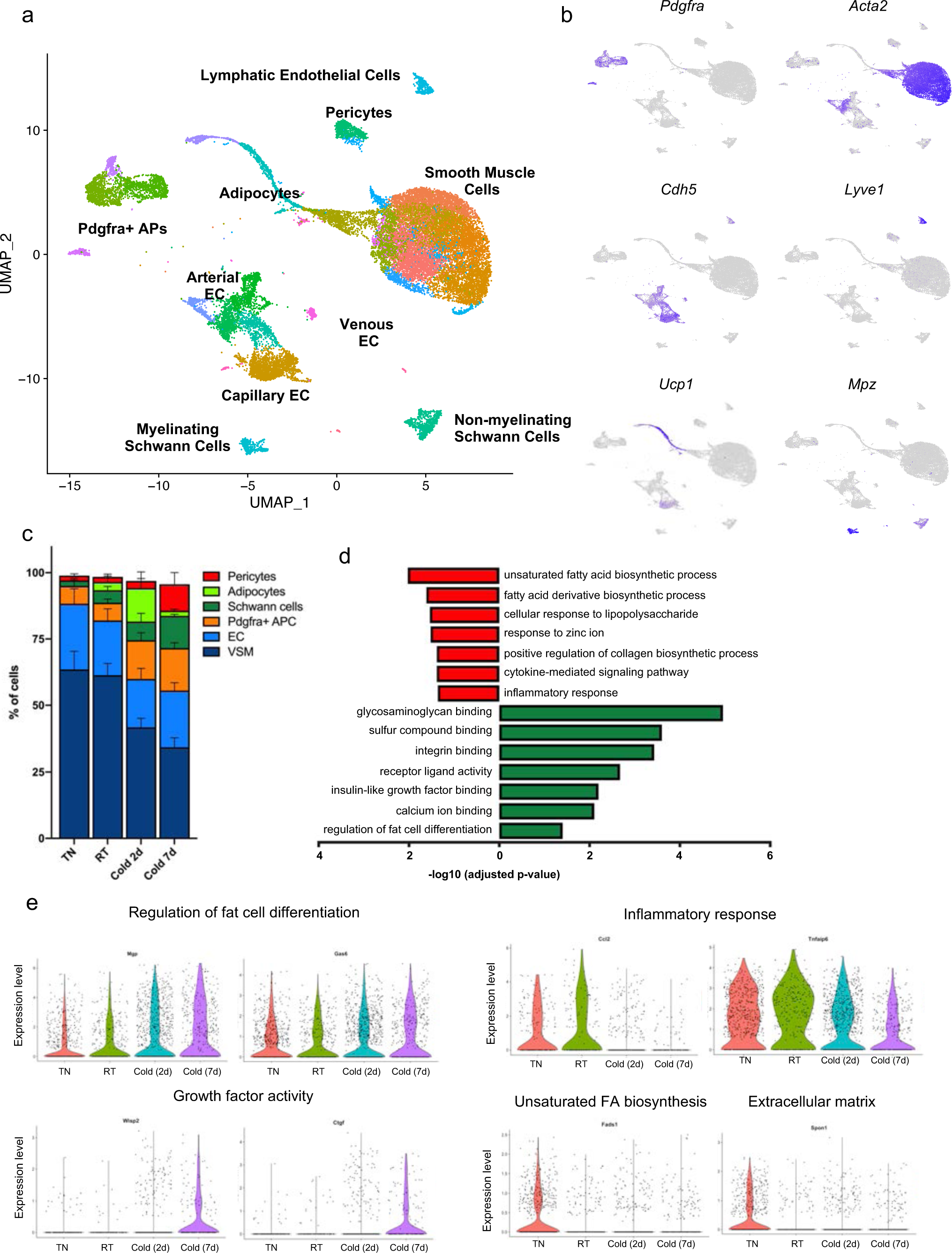
Single cell RNA-sequencing of the brown adipocyte niche during cold induced remodeling. (a) Unsupervised clustering of 24,498 non-immune cells from the BAT-SVF of 9-week old male C57BL/6J mice housed at TN (30 °C, 7 days), RT (22 °C), Cold (5 °C, 2 and 7 days) represented on a UMAP. (b) Individual gene UMAP plots showing the expression levels and distribution of representative marker genes. (c) Percentage of cells present in each cluster at different housing conditions. (d) Gene ontology (GO) enrichment analysis of transcripts upregulated (green) and downregulated (red) by cold in Pdgfra^pos^ adipocyte progenitor cells. (e) Violin plots of expression levels and distribution of representative transcripts for selected GO terms.

## Cold-induced remodeling of BAT involves cell-type specific events

Previous studies have shown that cold exposure activates *de novo* recruitment of brown adipocytes to increase maximal thermogenic activity^9,11,12^. To determine the cold-induced cellular dynamic within the brown adipose niche, we compared the frequency of different cell types identified in the scRNA-seq described above (Figure 1c). 7 days of cold exposure increased the percentage of Pdgfra^pos^ APC (p= 0.0453, Cold-7d vs TN) and Schwann cells (p= 0.0280, Cold-7d vs TN), while it reduced the percentage of VSMs (p< 0.0001, Cold-2d vs TN and Cold-7d vs TN). Interestingly, although we used a density-based fractionation of BAT to separate the mature lipid-laden adipocytes from the SVF, we could detect a number of cells expressing markers of brown adipocytes including *Ucp1, Adipoq, Cidea*, and *Dio2* (Figure 1b and Extended Data Figure 1). These cells were likely differentiating adipocytes with low lipid content and higher density than mature adipocytes. The frequency of these cells was significantly higher in the Cold-2d group (Figure 1c, p= 0.0124, Cold-2d vs TN), supporting the notion that cold exposure induces *de novo* adipogenesis in BAT.

The presence of differentiating adipocytes enabled us to explore the origin and adipogenic trajectory of brown adipocytes differentiated upon cold exposure. Given that previous studies have revealed the contribution of Pdgfra^pos^ Sca1^pos^ APC to cold-induced brown adipocyte differentiation^13^, we first examined the effect of cold exposure on gene expression in Pdgfra^pos^ APCs. To do so, we used a pseudo-bulk approach that requires aggregation of the single cell counts to the sample level for each cluster^21^ (Supplementary Table 2). Gene ontology analysis of the transcripts upregulated in Pdgfra^pos^ APCs by cold identified significant enrichments of glycosaminoglycan and integrin binding, insulin-like growth factor binding, and regulation of fat cell differentiation processes, all of which are known to be associated with adipogenesis^22,23^. The transcripts significantly downregulated by cold in Pdgfra^pos^ APCs were involved in unsaturated fatty acid biosynthesis, response to lipopolysaccharide, collagen biosynthesis, and inflammatory response processes (Figure 1d-e). Together, these analyses revealed that cold triggers a multi-layered gene program in Pdgfra^pos^ APC which support their adipogenic differentiation.

Using a similar strategy, we examined the effects of cold exposure on other cell types present in BAT niche (Supplementary Table 2 and Extended Data Figure 2-3. Description and discussion related to the ECs and Schwann cells are included in the respective figure legends). In VSMs, extracellular matrix organization, angiogenesis, cell division, cell junction assembly, epithelial cell migration, and response to TGFB stimulus were significantly over-represented in the transcripts whose expression was upregulated by cold (Extended Data Figure 4a-b). In the transcripts downregulated in cold, regulation of RNA splicing, ribosome biogenesis, striated muscle development, and G1/S transition of cell cycle were significantly over-represented (Extended Data Figure 4c-d).

## Lineage trajectory analysis predict the contribution of Trpv1^pos^ VSMs to brown adipogenesis

The UMAP cluster visualization showed the close relationship between differentiating brown adipocytes and VSMs (Figure 1a). This prompted us to interrogate the potential contribution of VSMs to the differentiating adipocytes using an *in silico* cell trajectory analysis method called Slingshot^24^, which is designed to infer cell lineages and model developmental trajectories from single-cell gene expression data. Consistent with Seurat clustering, pseudotemporal analysis predicted the VSMs to be the most likely origin of the differentiating brown adipocytes found in BAT-SVF (Figure 2a). To support these findings, we used another bioinformatics tool Vision to examine the coordinated expression of genes involved in known biological processes^25^. Consistently, we found high scores for adipogenesis and fatty acid metabolism gene signatures in VSMs (Extended Data Figure 5a), further highlighting their role as adipose precursors.

**Figure 2.**
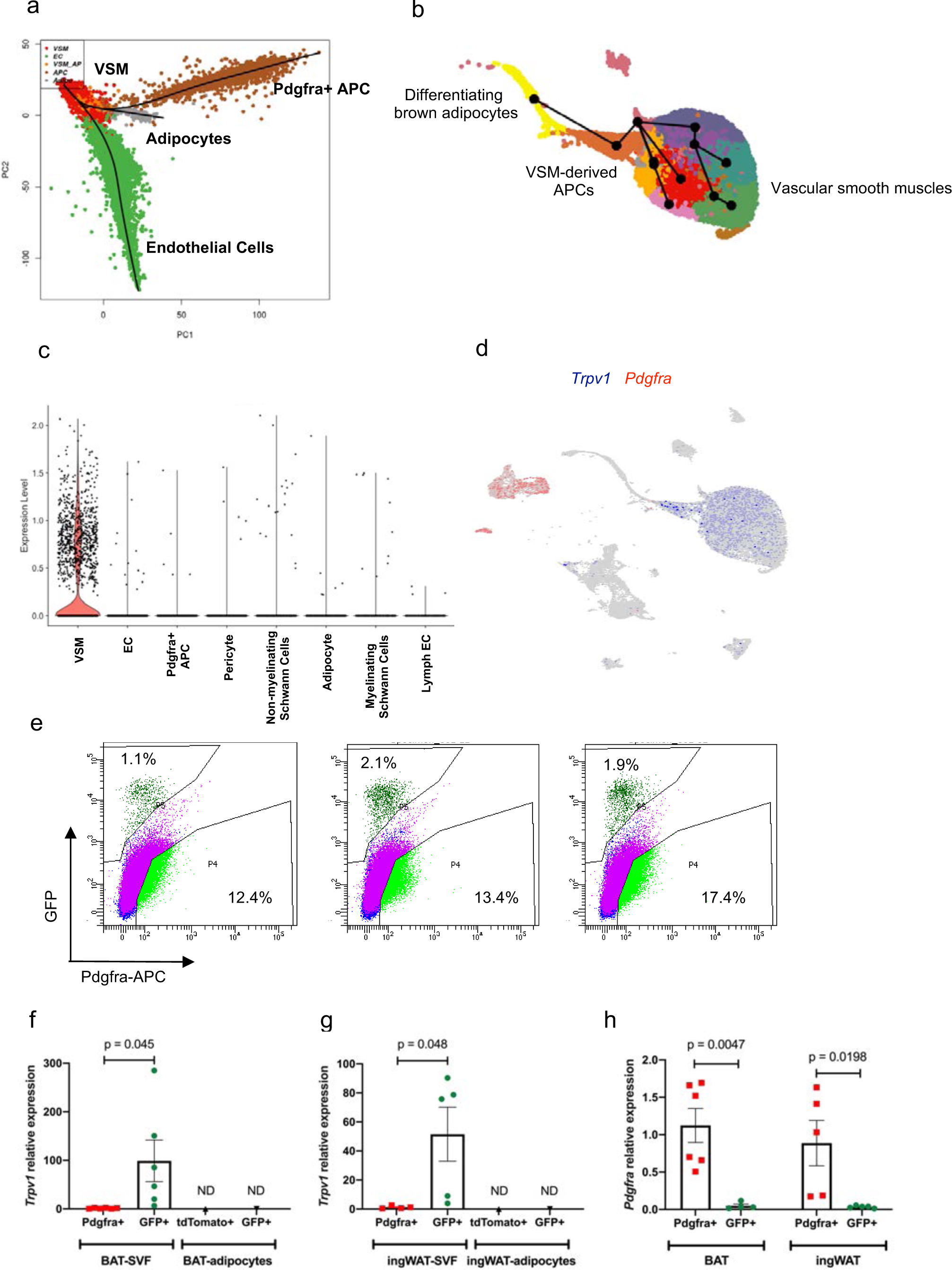
Lineage trajectory analysis predict the contribution of Trpv1^pos^ VSMs to brown adipogenesis. (a) Lineage trajectory analysis using Slingshot. Trajectories were calculated for the merged Seurat cell clusters, specifying the adipocyte cluster as the end state. (b) Pseudotime lineages calculated by Slingshot using the UMAP coordinates and cell clusters for the vascular smooth muscle cells and adipocytes. (c) Violin plots showing *Trpv1* expression in different cell types present in BAT. (d) UMAP plot showing expression of *Trpv1* and *Pdgfr*a in BAT-SVF. (e) Representative flow cytometry analysis of BAT-SVF derived from Trpv1^cre^ Rosa26^mTmG^ mice stained with Pdgfra-APC. Trpv1 expression in Pdgfra^pos^ and GFP^pos^ cells from (f) BAT-SVF and (g) ingWAT-SVF, in addition to tdTomato^pos^ and GFP^pos^ adipocytes from each depot. N=6 per group. Data are presented as means ± SEM, with significance tested by unpaired T-test in e and f. (h) *Pdgfr*a expression in Pdgfra^pos^ and GFP^pos^ cells from BAT-SVF and ingWAT-SVF. N=6 per group. Significance was tested by Two-way ANOVA with Sidak’s multiple comparisons test.

Although several studies have demonstrated that VSM-derived APCs in white adipose tissue can differentiate into white or beige adipocytes^15,16,26-28^, lineage tracing experiments demonstrate that the Myh11-expressing^16^ or SMA-expressing^15^ VSM cells do not give rise to the classical brown adipocytes. To identify the sub-populations of VSMs with brown adipogenic potential and predict the adipogenic trajectory, we used Slingshot trajectory analysis and illustrated the progression of VSM cells to adipocyte progenitors and differentiating adipocytes (Figure 2b). This notion was further supported by pathway enrichment analysis of the top 100 genes correlated with the adipogenic trajectory (pseudotime genes), which showed the enrichment of thermogenic adipocyte features, such as oxidative phosphorylation, heat production by uncoupling protein, TCA cycle, PPAR signaling, and triglyceride biosynthesis pathways (Extended Data Figure 5b).

To pinpoint a specific marker to track the fate of VSMs in BAT, we identified *Trpv1* as one of the most exclusively expressed transcripts in the VSM cells with high expression in the predicted VSM-derived APCs (Figure 2c-d). The scRNA-seq data suggested that Trpv1^pos^ VSMs are a distinct population of cells in BAT and they do not have any overlap with the Pdgfra^pos^ APCs (Figure 2d).

To validate these findings, we generated a Trpv1 reporter mouse model by crossing the Trpv1-cre mice with the Rosa26-mTmG reporter strain (Trpv1^Cre^ Rosa26^mTmG^). In the absence of cre recombinase, the membranes of all the cells are labeled with tdTomato. Cells that express Trpv1 undergo cre recombination and their membranes are permanently labeled with GFP. Flow cytometry analysis of cells in the BAT-SVF of Trpv1^Cre^ Rosa26^mTmG^ revealed that GFP labeling and Pdgfra-expression are mutually exclusive, and the GFP-labeled cells are in fact a distinct population of cells in BAT from putative Pdgfra^pos^ adipocyte progenitor cells (Figure 2e and Extended Data Figure 6). Gene expression analysis in the isolated cells also confirmed that *Trpv1* and *Pdgfra* expression are enriched in the GFP^pos^ and Pdgfra^pos^ cells isolated from the SVF, respectively (Figure 2f-h). Additionally, similar to *Pdgfra, Trpv1* expression is restricted to the progenitor cell populations in the SVF of BAT and inguinal white adipose tissue (ingWAT) and is not detected in mature brown or white adipocytes, consistent with multiple reports that TRPV1 expression decreases during adipogenesis^29,30^ (Figure 2f-g).

## Trpv1-expressing VSM-APCs give rise to brown and white adipocytes

The lineage trajectory analysis predicted that Trpv1-expressing VSMs are a cellular origin for differentiating brown adipocytes. To test this hypothesis and examine the adipogenic potential of Trpv1-expressing cells *in vivo*, we performed lineage tracing studies using the Trpv1^Cre^ Rosa26^mTmG^ mouse model (Figure 3a). Given the absence of Trpv1 expression in adipocytes and the indelible nature of the GFP labeling, we reasoned that the presence of GFP-labeled adipocytes would mean that Trpv1^pos^ VSMs could differentiate into adipocytes. Examining GFP and tdTomato fluorescence in BAT of male and female Trpv1^Cre^ Rosa26^mTmG^ mice housed at room temperature demonstrated the presence of GFP-labeled adipocytes in BAT (Figure 3b). Immunohistochemistry using GFP and perilipin (Plin1) antibodies confirmed the colocalization of GFP and Plin1 on the same cells, confirming that the GFP-labeled cells are in fact mature adipocytes (Figure 3c). Interestingly, we found that Trpv1^pos^ progenitors also contribute to the white adipocytes in ingWAT and perigonadal WAT (pgWAT), as evidenced by the presence of GFP-labeled adipocytes in both WAT depots of the Trpv1^Cre^ Rosa26^mTmG^ mice (Figure 3b). Together, these results not only validated the findings of the scRNA-seq, but also identified a population of APCs derived from the Trpv1^pos^-VSMs in brown and white adipose tissue which are capable of generating mature adipocytes.

**Figure 3.**
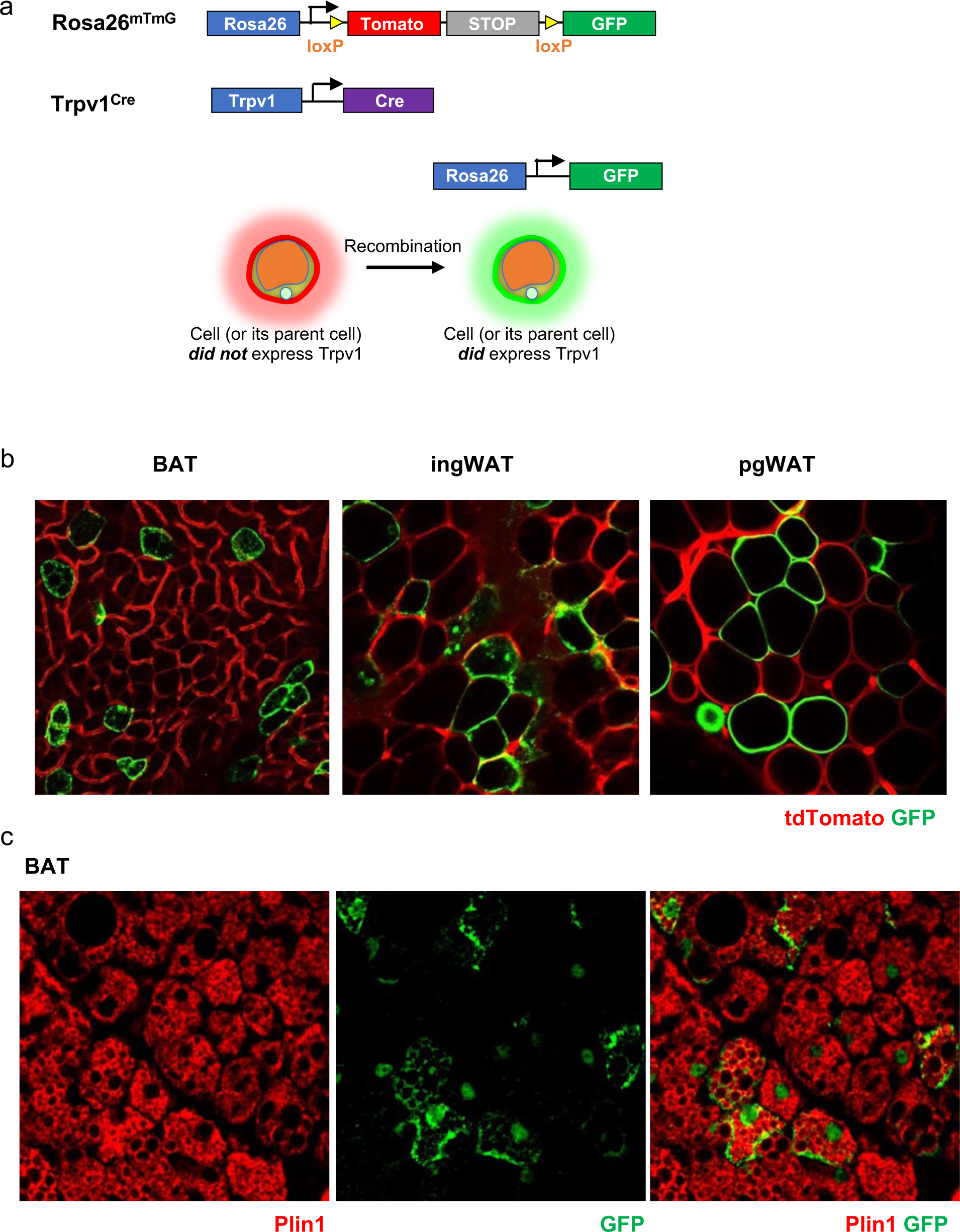
Genetic lineage tracing shows that Trpv1^pos^ cells give rise to brown and white adipocytes. (a) Scheme of Trpv1^cre^ Rosa26^mTmG^ lineage tracing model. (b) Whole-mount microscopy for GFP and tdTomato fluorescence in BAT, ingWAT, and pgWAT of Trpv1^cre^ Rosa26^mTmG^ mice.

## Cold exposure induces the differentiation of Trpv1^pos^ progenitors into highly thermogenic adipocytes

Although cold exposure (2 days or 7 days) did not change the expression of Trpv1 in the VSMs of BAT (Figure 4a), we observed a significant increase in the number of differentiating adipocytes in the BAT of mice housed in cold for 2 days (Figure 4b). This prompted us to investigate if cold exposure increases the number of brown adipocytes derived from the Trpv1^pos^ lineage. To test this hypothesis, we housed Trpv1^Cre^ Rosa26^mTmG^ mice at 5 °C for 7 days and examined GFP and tdTomato fluorescence in BAT, ingWAT, and pgWAT compared to the control mice kept at RT. Quantification of fluorescence intensity revealed that cold exposure significantly increased the number of GFP+ adipocytes in BAT (Figure 4c), but not in ingWAT or pgWAT (Extended Data Figure 7a-b).

**Figure 4.**
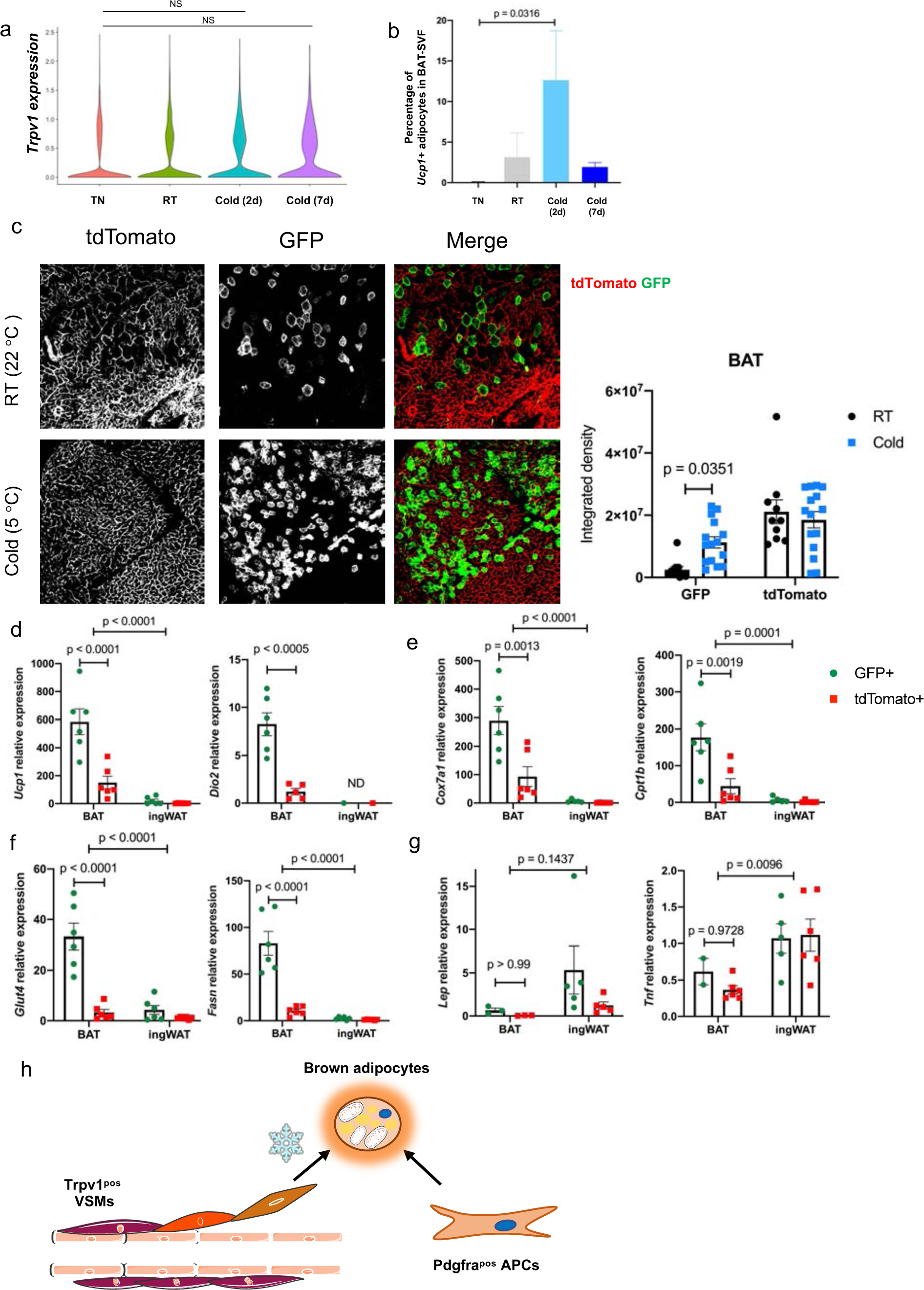
Cold exposure induces differentiation of Trpv1^pos^ adipocyte progenitors into thermogenic adipocytes. (a) Violin plot showing *Trpv1* expression in BAT-SVF at different housing conditions. (b) Percentage of cells in brown adipocyte cluster at different housing conditions. N=4 per group. Data are presented as Means ± SEM with significance tested by One-Way ANOVA with Dunnett’s multiple comparisons test. (c) Left: Whole-mount microscopy of GFP and tdTomato fluorescence in BAT of Trpv1^cre^ Rosa26^mTmG^ mice housed at RT or Cold for 7 days. Right: Quantification of GFP and tdTomato signal (integrated density) in each depot. N=3 mice per group, 3-8 slides/mouse. Data are presented as Means ± SEM with significance tested by Two-Way ANOVA with Sidak’s multiple comparisons test. (d-g) Expression of key adipogenic genes in tdTomato^pos^ and GFP^pos^ adipocytes isolated from BAT and ingWAT of Trpv1^cre^ Rosa26^mTmG^ housed in cold (5 °C) for 7 days. N=6 mice per group. Data are presented as Means ± SEM with significance tested by Two-Way ANOVA with Sidak’s multiple comparisons test. (h) Model depicting the two types of brown adipocyte progenitors contributing to the mature brown adipocyte pool in adult mice.

These findings demonstrated that brown adipocytes could arise from both Trpv1^pos^ and Trpv1^neg^ lineages, but cold exposure specifically induced differentiation of Trpv1^pos^ progenitors. To determine whether brown adipocytes derived from either lineage possess intrinsic differences, we isolated GFP^pos^ (derived from Trpv1^pos^ APCs) and tdTomato^pos^ (derived from Trpv1^neg^ APCs) mature adipocytes from BAT and ingWAT of the Trpv1^Cre^ Rosa26^mTmG^ mice and measured gene expression. Intriguingly, compared to the tdTomato^pos^ adipocytes, the GFP^pos^ brown adipocytes isolated from BAT expressed significantly higher levels of the key thermogenic genes (*Ucp1, Cidea, Dio2, Ppargc1a*) (Figure 4d and Extended Data Figure 7c), mitochondrial genes (*Cox5b, Cox7a1, Cpt1b*) (Figure 4e and Extended Data Figure 7d), as well as other genes regulating fuel uptake and utilization (*Glut4, Gapdh*, and *Fasn*) (Figure 4f and Extended Data Figure 7e). On the contrary, the expression of most of these genes was similar between the GFP^pos^ and tdTomato^pos^ adipocytes isolated from ingWAT. Levels of general adipogenic transcription factor *Pparg* was comparable between adipocytes isolated from BAT and ingWAT (Extended Data Figure 7e). Additionally, the expression of white adipocyte-enriched genes, *Lep, Retn*, and *Tnf* was higher in the adipocytes isolated from ingWAT than those isolated from BAT and was not different between the GFP^pos^ and tdTomato^pos^ adipocytes (Figure 4g and Extended Data Figure 7f). Together, these results show that cold exposure promotes the adipogenic differentiation of Trpv1^pos^ VSMs in BAT, which give rise to brown adipocytes with high thermogenic potential.

Searching for potential underlying mechanisms for VSM cells to acquire adipogenic potential, we found a gene signature of Epithelial to Mesenchymal Transition (EMT), driven by differences in key genes *Snai2, Zeb2*, and *Tcf4* as well as *Cdh2*^*31*^ was associated with the VSM transition to adipocytes (Extended Data Figure 8). EMT involves major remodeling of cell-cell and cell-extracellular matrix interactions which allow migration and transition of epithelial cells to the mesenchymal fate^32^. Indeed, we found cold exposure induced the expression of genes involved in EMT-related pathways including cell junction assembly and epithelial cell migration in VSMs (Extended Data Figure 4a-b), suggesting that cold exposure triggers EMT in the VSMs of BAT, which may facilitate their contribution to brown adipogenesis.

Recent technical advances allowing single cell approaches, combined with development of powerful computation analyses, have provided an opportunity to address fundamental questions in adipocyte biology^33,34^. We use scRNA-seq to map the cellular composition of BAT during cold exposure and identify the cellular origin of brown adipocytes recruited in BAT upon cold challenge. In addition to previously known APCs, we identified the VSM-derived Trpv1^pos^ cells as a novel population of adipocyte progenitors. Genetic lineage tracing of the Trpv1^pos^ progenitors showed contribution of these cells to adipocytes in BAT, ingWAT, and pgWAT depots. Based on previous knowledge of the contribution of Pdgfra^pos^ mesenchymal progenitors to brown adipocytes^13^ and our new findings, we propose a model in which cold stimulates both Trpv1^pos^ APCs and Pdgfra^pos^ APCs to undergo adipogenesis and expand the brown adipocyte pool to adapt to increased thermogenic demand. Cold exposure induces the expression of transcripts involved in epithelial cell migration and cell junction assembly in Trpv1^pos^ APCs and facilitates their differentiation to highly thermogenic adipocytes (Figure 4h). Importantly, we show that brown adipocytes derived from the Trpv1^pos^ lineage express higher levels of thermogenic genes compared to those derived from the Trpv1^neg^ progenitors.

Lineage tracing studies using multiple Cre drivers labeling the VSM lineage have tracked the differentiation of perivascular smooth muscle cells to the white and beige adipocyte pool^15,16,26,28^. Disrupting adipogenic differentiation of selective VSM progenitors blunts beige adipocyte formation and impairs thermoregulation as well as glycemic control in mice^15,26^. However, fate-mapping studies have failed to show that brown adipocytes are derived from those selective VSM lineage in interscapular BAT^15,16^. The major limitation of these studies is that they have relied on a small number of common VSM lineage markers to assess adipogenic potential and track their transition to mature adipocyte fate. Our scRNA-seq data clearly point out the heterogeneity of vascular smooth muscle cells within the brown adipose niche. In contrast to the previous studies, we used an unbiased strategy to identify the origin of the cold-induced brown adipocytes, which uncovered Trpv1-expressing VSMs as progenitors giving rise to adipocytes in BAT, ingWAT, and pgWAT depots.

Trpv1 is a cell surface receptor that detects multiple noxious stimuli including heat, pH changes, fatty acid amides, and capsaicin (the pungent component of chilli-peppers)^35^. In addition to sensory neurons, Trpv1 is expressed in smooth muscle cells of several thermoregulatory tissues, including the skin, ear, cremaster muscle, dura, tongue, and trachea^36^. In this study, we report Trpv1-expressing VSMs in the major thermogenic organ BAT. Emerging evidence suggest a role for TRPV1 signaling in energy metabolism and feeding. Administration of TRPV1 agonist capsaicin and its related compounds reduces food intake and increases energy expenditure in animals and humans^29,37-39^. In mice, capsaicin activates recruitment of beige adipocytes in a TRPV1 dependent manner^40^, however curiously TRPV1-deficient mice have higher energy expenditure and are protected from diet-induced obesity^41^. Additionally, capsaicin treatment blocks adipogenesis of 3T3-L1-preadipocytes in a Trpv1-dependent manner^29^. Here we show that cold stimulates the differentiation of Trpv1^pos^ adipocyte progenitors to mature adipocytes. The identification of Trpv1^pos^ adipocyte progenitors and their role in cold-induced brown adipocyte recruitment will enable future investigations to address the role of Trpv1 channel in thermoregulation.

In summary, these findings suggest a new model for the development of BAT that could be critical in designing strategies to increase the number of thermogenic adipocytes as a therapeutic approach for obesity and metabolic diseases.

---

**Extended Data Figure 1.**
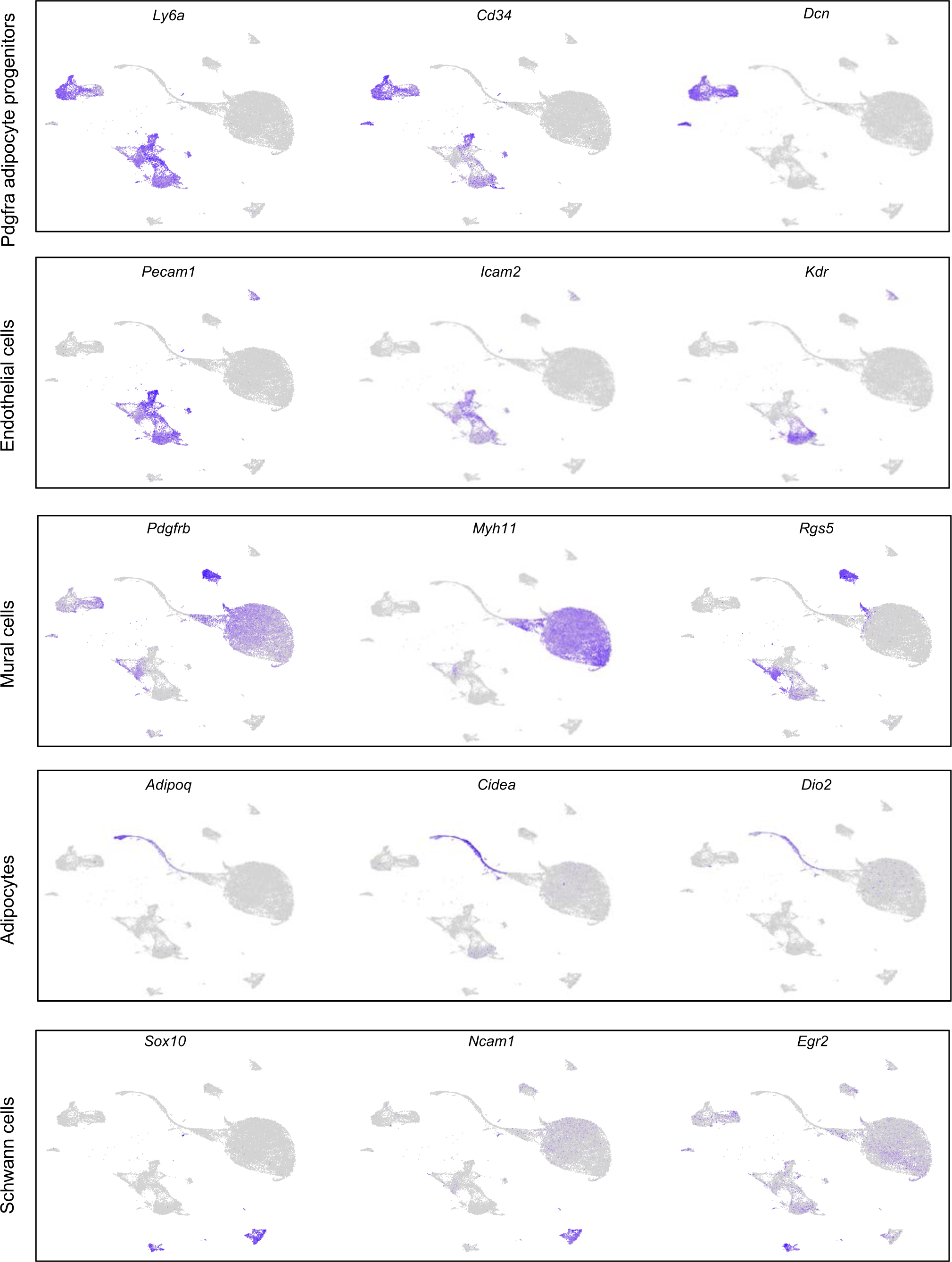
Characterization of cell types present in the BAT-SVF, related to Figure 1. Individual gene UMAP plots showing the expression levels and distribution of representative marker genes for each cell type.

**Extended Data Figure 2.**
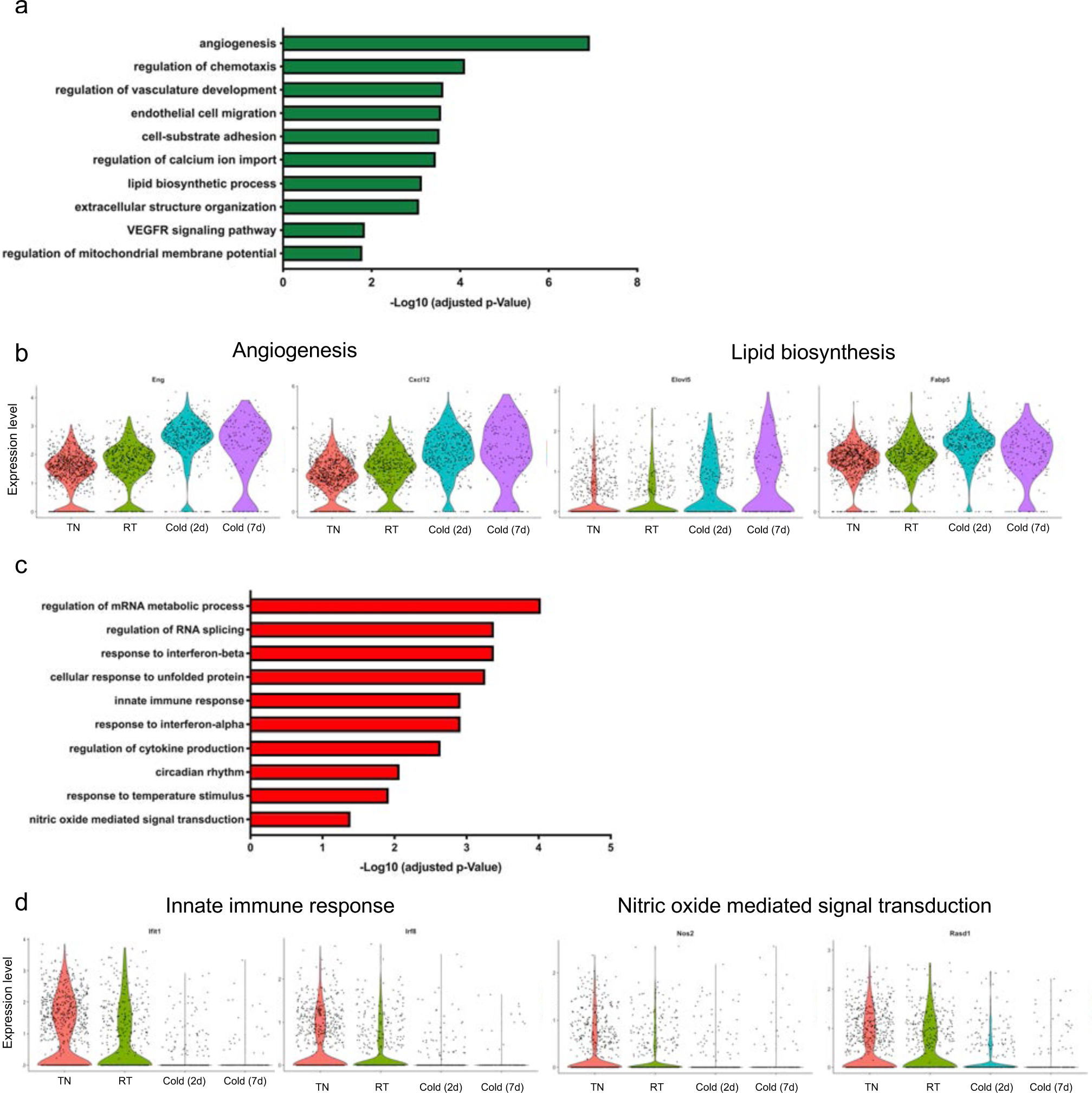
Cold exposure triggers the remodeling and expansion of endothelial cells, related to Figure 1. (a) Gene ontology (GO) enrichment analysis of the transcripts upregulated in capillary endothelial cells by cold. (b) Violin plots showing the expression levels and distribution of representative upregulated transcripts for the selected GO terms. (c) Gene ontology (GO) enrichment analysis of the transcripts downregulated in capillary endothelial cells by cold. (d) Violin plots showing the expression levels and distribution of representative downregulated transcripts for the selected GO terms. In capillary endothelial cells, cold exposure upregulated the expression of transcripts involved in angiogenesis, chemotaxis, endothelial cell migration, and lipid biosynthesis (Extended Data Figure 2a-b). This is consistent with previous studies reporting increased angiogenesis and expansion of vascular networks in BAT upon cold exposure^42^. On the other hand, cold exposure downregulated the expression of genes related to mRNA metabolism, response to unfolded protein, innate immune response, cytokine production, and Nitric oxide mediated signal transduction in capillary endothelial cells (Extended Data Figure 2c-d).

**Extended Data Figure 3.**
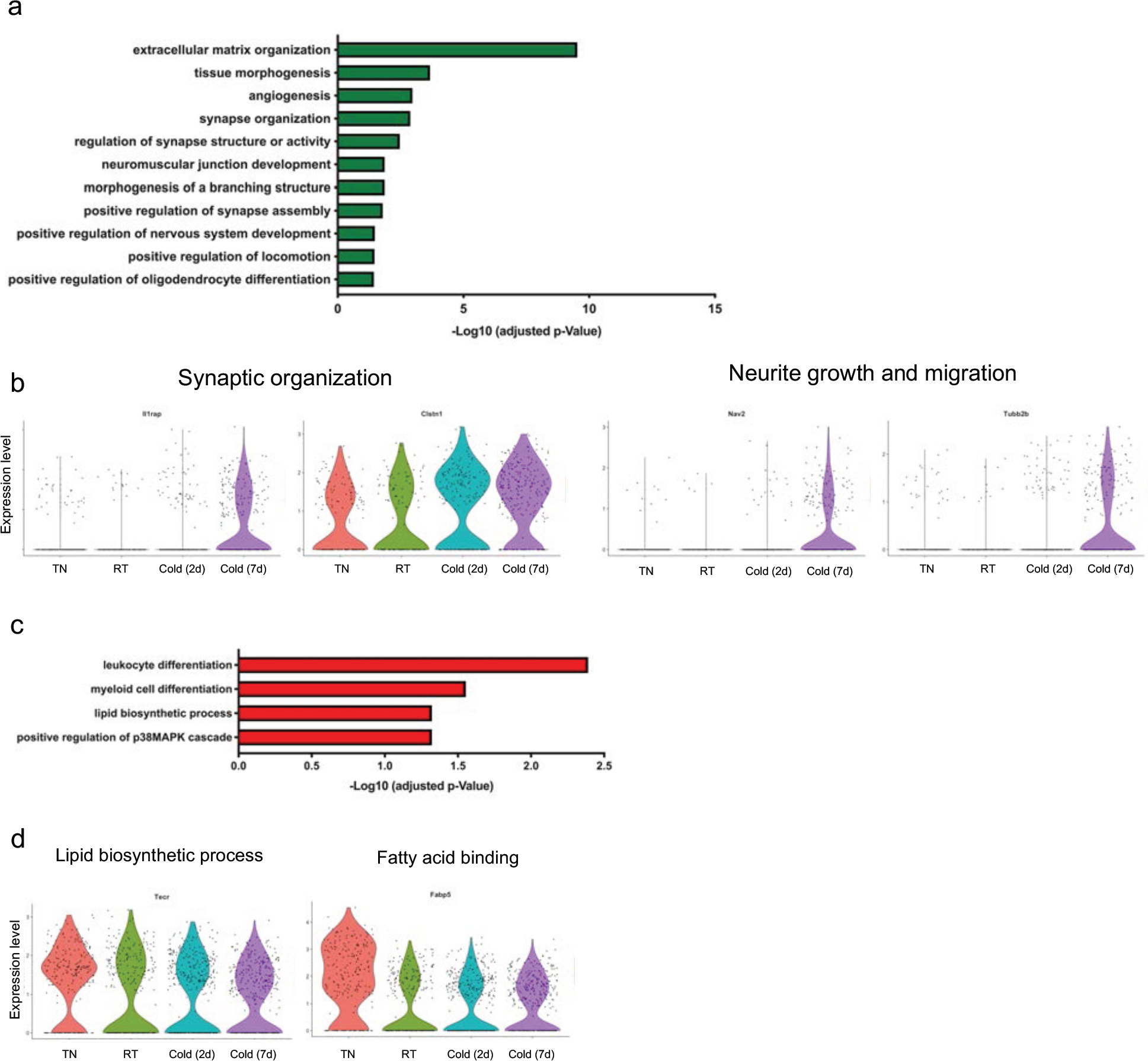
Cold exposure promotes the growth and reorganization of Schwann cells, related to Figure 1. (a) Gene ontology (GO) enrichment analysis of the transcripts upregulated in Schwann cells by cold. (b) Violin plots showing the expression levels and distribution of representative upregulated transcripts for the selected GO terms. (c) Gene ontology (GO) enrichment analysis of the transcripts downregulated in Schwann cells by cold. (d) Violin plots showing the expression levels and distribution of representative downregulated transcripts for the selected GO terms. In Schwann cells, the transcripts upregulated by cold belong to biological processes including extracellular matrix organization, regulation of synapse structure or activity, positive regulation of synapse assembly, and oligodendrocyte differentiation (Extended Data Figure 3a-b). The activation of these pathways enables the appropriate expansion of Schwann cells and sympathetic neurites to promote norepinephrine-mediated activation of brown adipocytes by cold^43^. The transcripts downregulated by cold were involved in leukocyte differentiation, lipid biosynthesis, and positive regulation of p38MAPK cascade (Extended Data Figure 3c-d).

**Extended Data Figure 4.**
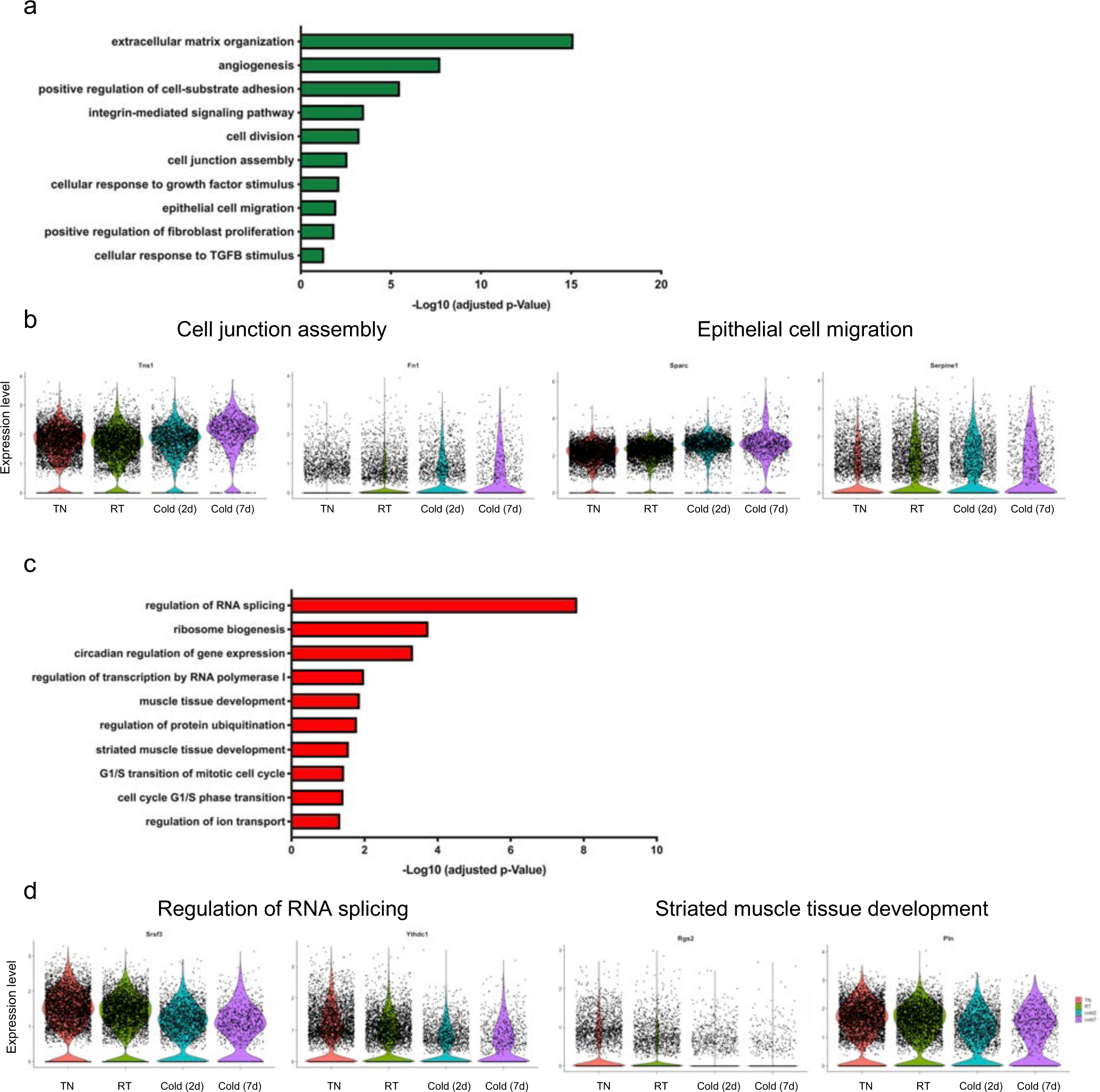
Cold-induced transcriptional changes in vascular smooth muscles, related to Figure 1. (a) Gene ontology (GO) enrichment analysis of the transcripts upregulated in vascular smooth muscles by cold. (b) Violin plots showing the expression levels and distribution of representative upregulated transcripts for the selected GO terms. (c) Gene ontology (GO) enrichment analysis of the transcripts downregulated in vascular smooth muscles by cold. (d) Violin plots showing the expression levels and distribution of representative downregulated transcripts for the selected GO terms.

**Extended Data Figure 5.**
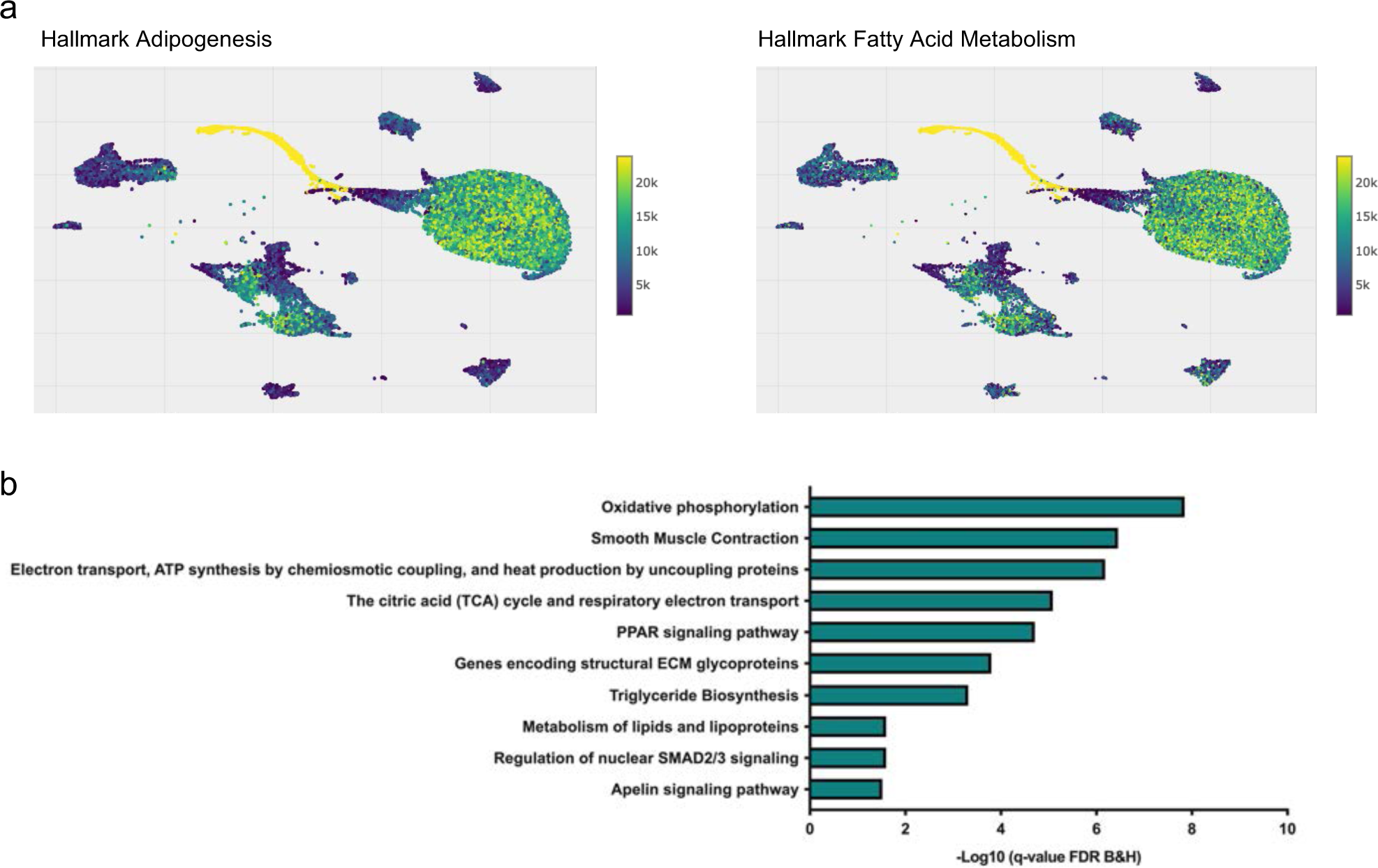
Modeling the differentiation trajectory of VSM-derived progenitors to adipocytes, related to Figure 2. (a) Adipogenesis and Fatty acid metabolism gene signatures visualized on UMAP plots using VISION. (b) Pathway enrichment analysis of top 100 genes driving the trajectory.

**Extended Data Figure 6.**
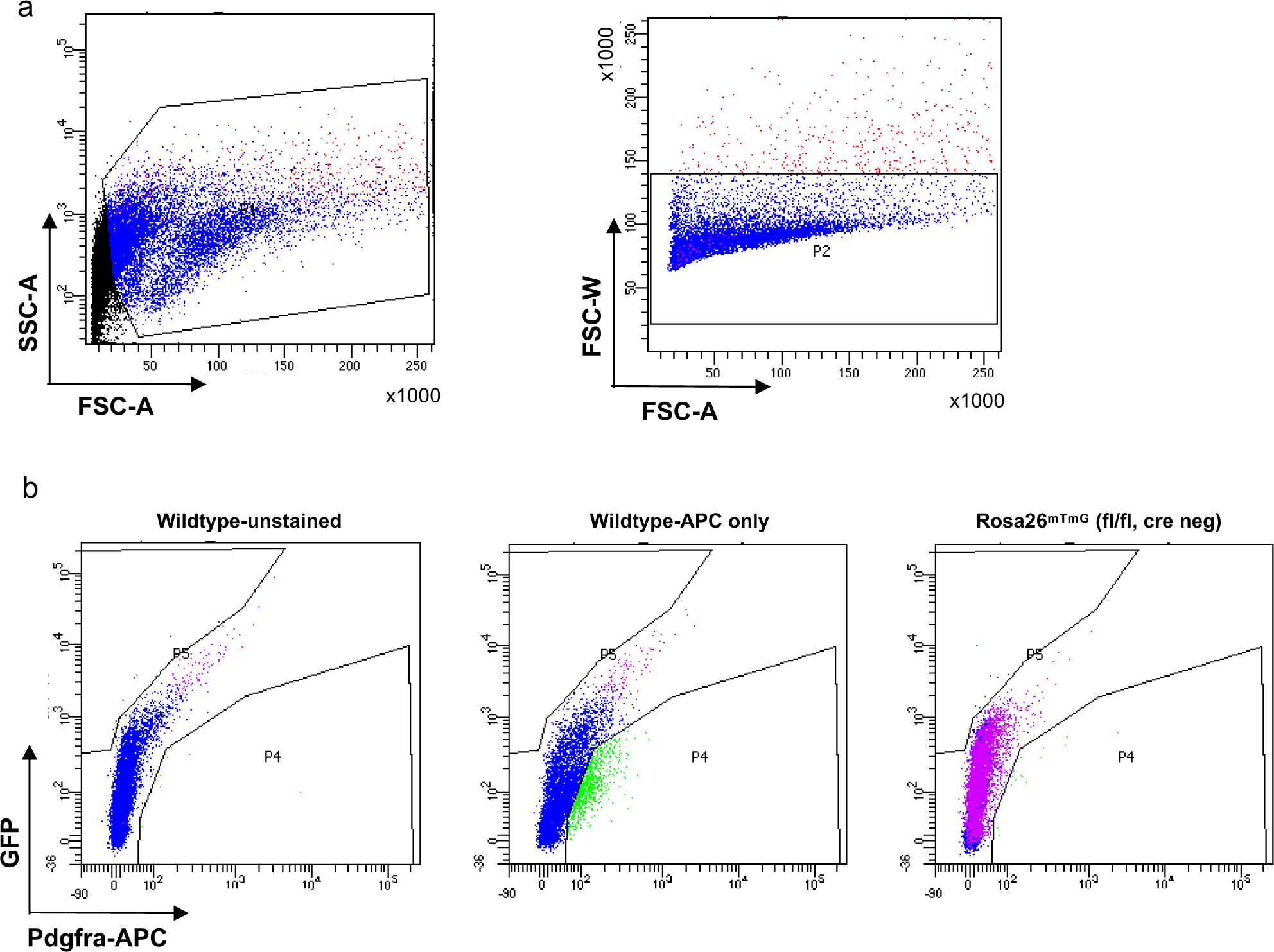
Gating strategy in flow cytometry analysis in BAT-SVF of Trpv1^cre^ Rosa26^mTmG^, related to Figure 2. (a) Debris and doublets were excluded based on forward and side scatter profiles. (b) GFP and APC labeling in samples with neither APC or GFP labeling (Wildtype-unstained), APC labeling (Wildtype-APC only), and tdTomato labeling (Rosa26^mTmG^ (fl/fl, cre neg)).

**Extended Data Figure 7.**
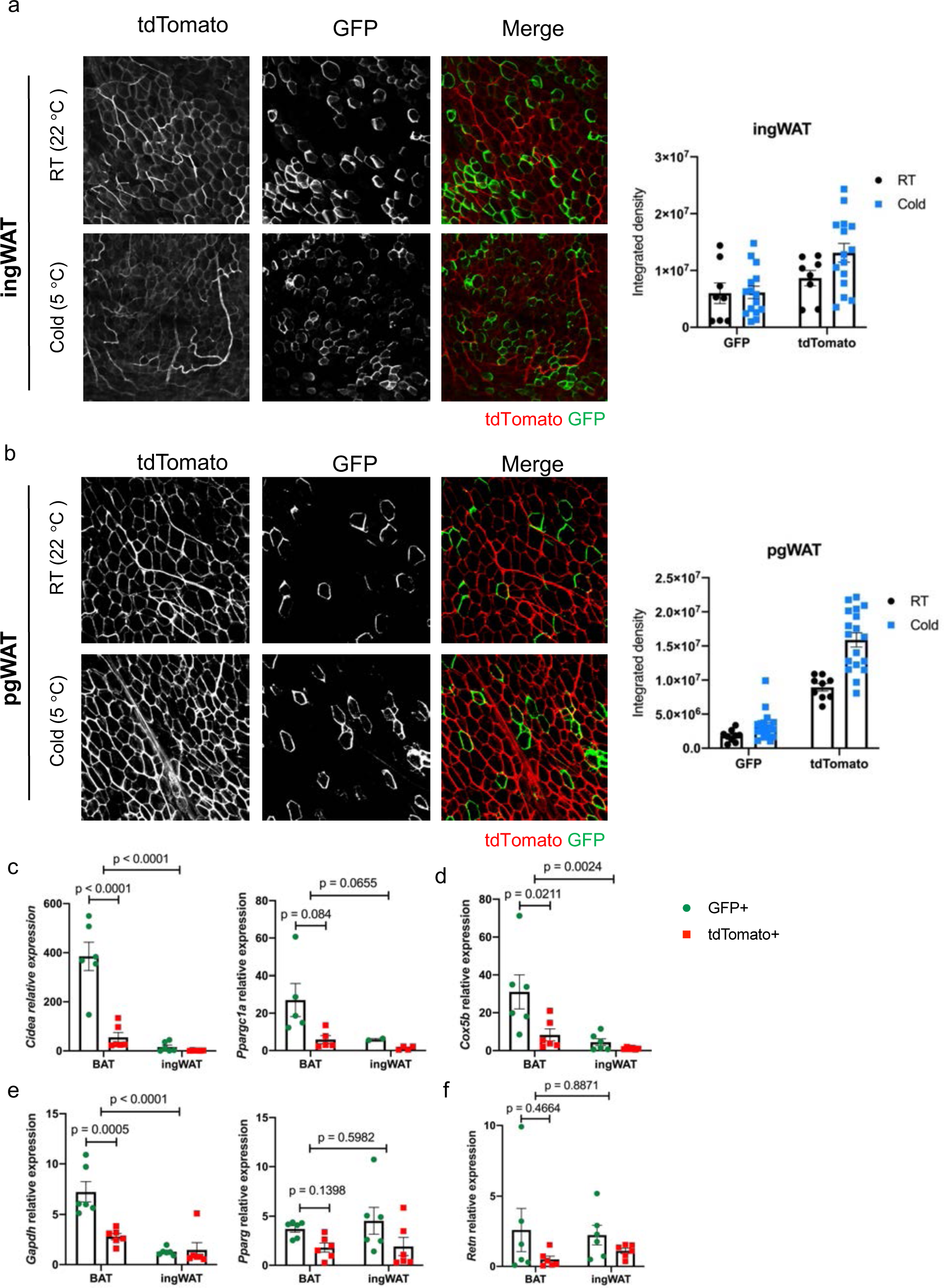
Cold does not increase the number of GFP^pos^ adipocytes in WAT, related to Figure 4. Left: Whole-mount microscopy for GFP and tdTomato fluorescence in (a) ingWAT and (b) pgWAT of Trpv1^cre^ Rosa26^mTmG^ mice housed at RT or cold for 7 days. Right: Quantification of GFP and tdTomato signal (integrated density) in each depot. N=3 mice per group, 3-8 slides/mouse. Data are presented as Means ± SEM. (c-f) Expression of the indicated transcripts in tdTomato^pos^ and GFP^pos^ adipocytes isolated from BAT and ingWAT of Trpv1^cre^ Rosa26^mTmG^ housed at Cold (5 °C) for 7 days. N=6 mice per group. Data are presented as Means ± SEM. Two-Way ANOVA with Sidak’s multiple comparisons test.

**Extended Data Figure 8.**
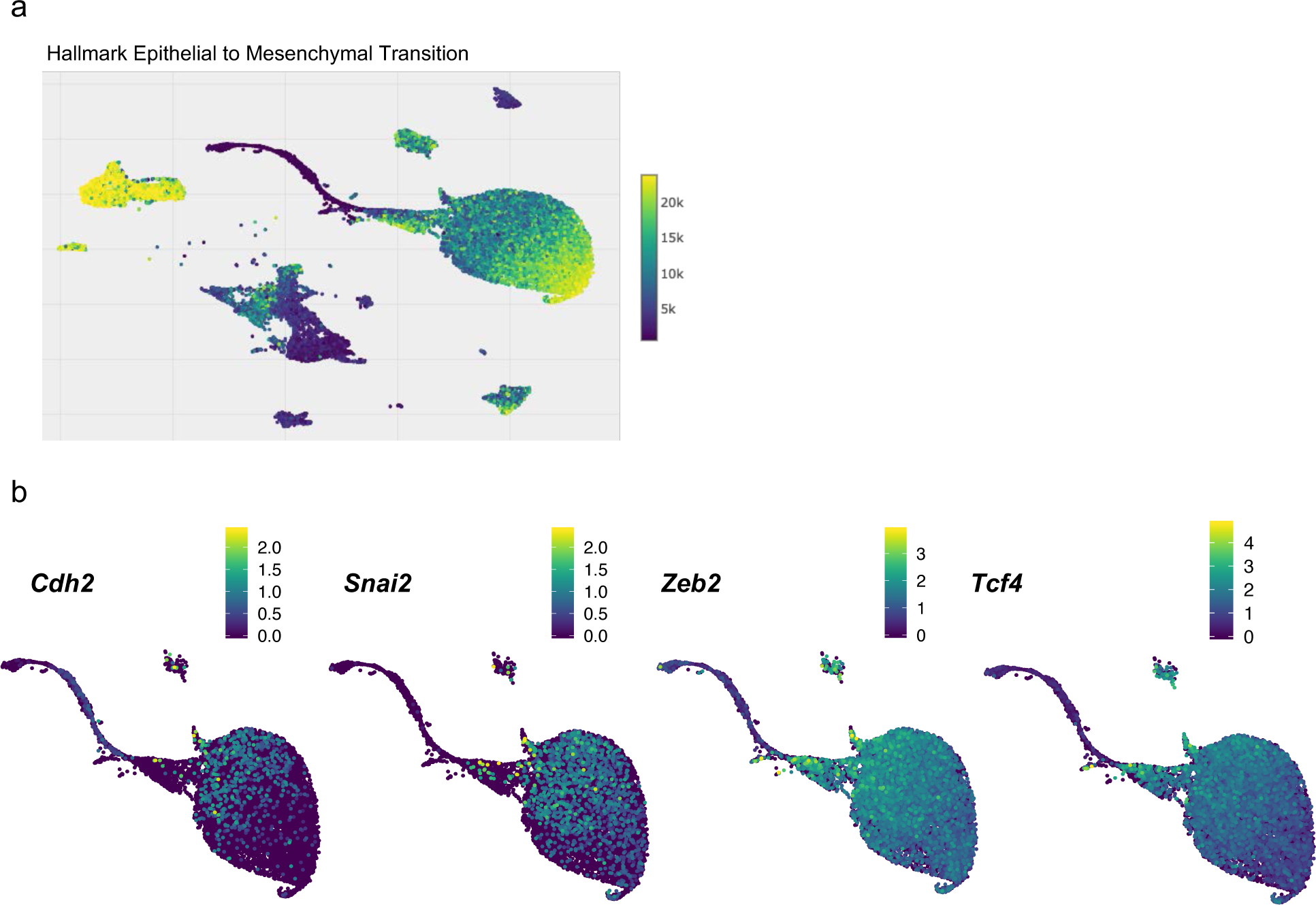
VSMs undergo Epithelial to Mesenchymal Transition, related to Figure 4. (a) Epithelial to Mesenchymal Transition gene signature visualized on UMAP plots using VISION. (b) Expression of the transcripts involved in epithelial to mesenchymal transition visualized on UMAP plots.

**Extended Data Table 1.**
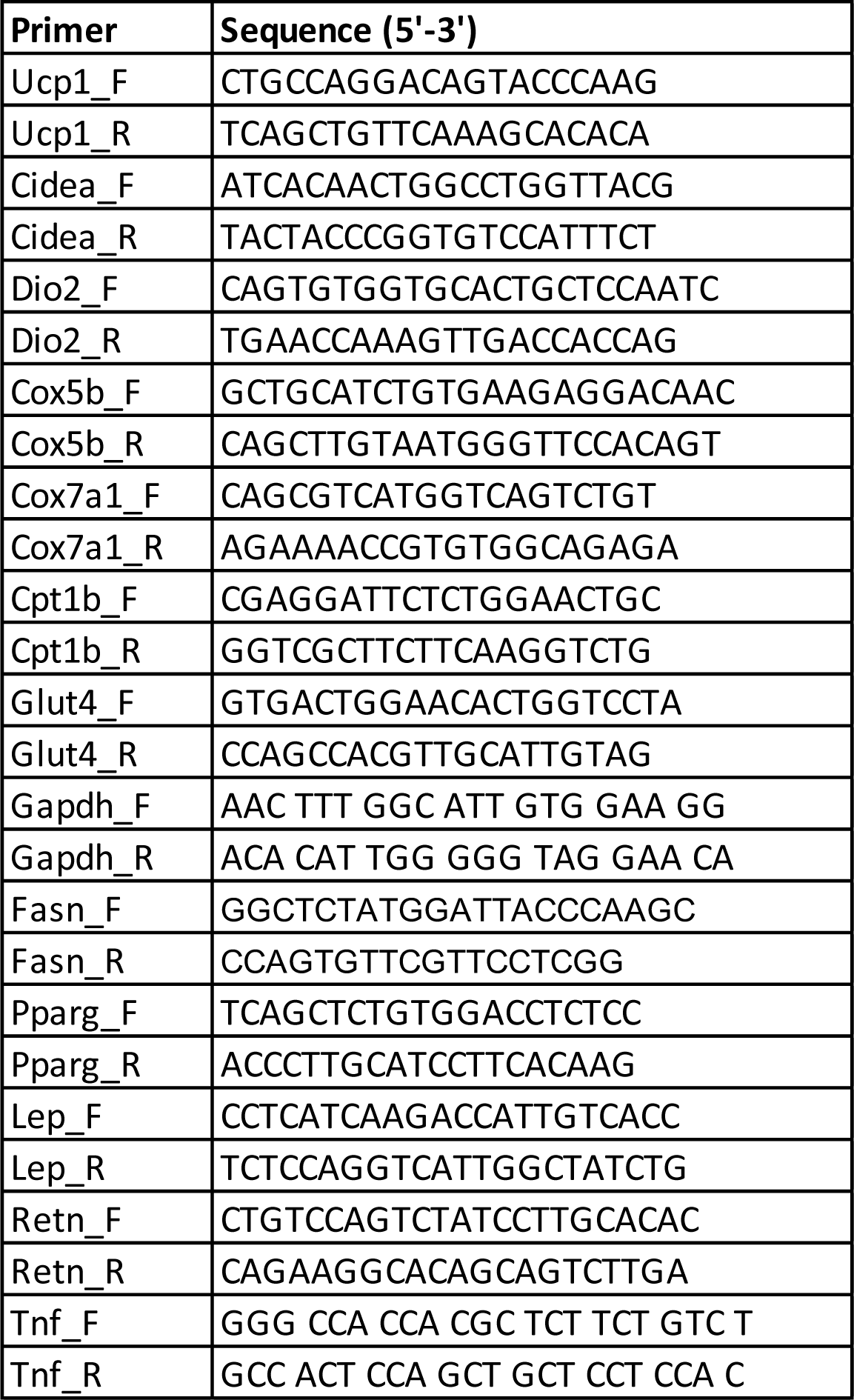
qPCR primer sequences.

## Methods

### Animals

Study approval: All experimental procedures involving animals were performed in compliance with relevant ethical regulations applied to the use of small rodents, and with approval by the Institutional Animal Care and Use Committees (IACUC) at Joslin Diabetes Center.

C57Bl/6J mice (Stock no. 000664) were purchased from The Jackson Laboratory. For experiments involving cold exposure, C57BL6J mice were housed at 5°C (cold) or 30°C (thermoneutral) in a controlled environmental diurnal chamber (Caron Products & Services Inc., Marietta, OH) with free access to food and water.

Generation of Rosa26-mTmG reporter strain: Homozygous Trpv1^cre^ mice (B6.129-Trpv1tm1(cre)Bbm/J, Stock No: 017769) were crossed with Homozygous Rosa26^mTmG^ (B6.129(Cg)-Gt(ROSA)26Sortm4(ACTB-tdTomato,-EGFP)Luo/J, Stock no. 007676), both obtained from The Jackson Laboratory.

### Isolation of stromovascular fraction from BAT

Interscapular BAT was dissected, minced, and digested with a cocktail containing type 1 Collagenase (1.5 mg/mL; Worthington Biochemical), Dispase II (2.5 U/mL; STEMCELL Technologies), fatty acid free bovine serum albumin (2%; Gemini Bio Products) in Hanks’ balanced salt’s solution (Corning® Hank’s Balanced Salt Solution, 1X with calcium and magnesium) for 45 minutes at 37°C with gentle shaking. Dissociated tissue was centrifuged at 500 g at 4°C for 10 minutes. The top adipocyte layer and the supernatant were gently removed, the SVF pellet was resuspended in 10 ml of 10% Fetal Bovine Serum (FBS) in DMEM, filtered through a 100 µm cell strainer into a fresh 50 ml tube, and centrifuged at 500 g for 7 minutes. The red blood cell lysis was performed by resuspending the pellet in 2 ml sterile ACK (Ammonium-Chloride-Potassium) lysis buffer (ACK Lysing Buffer, Lonza) and incubating on ice for 5 minutes. The cells were then filtered through a 40 µm cell strainer, washed with 20 ml 10% FBS in DMEM, and centrifuged at 500 g for 7 minutes. The pellet was resuspended in 1 ml of 1.5% BSA in PBS. The dead cell removal was performed using Dead Cell Removal Kit (Miltenyi Biotec) according to the manufacturer’s instructions. The cells were finally resuspended in 50-100 µl of 1.5% BSA in PBS and kept on ice before immediately proceeding to single cell isolation.

### Single-cell RNA-sequencing

Cells were loaded for an expected recovery of 10,000 cells per channel. The loaded Chip with single cell suspension was placed on a 10x Genomics Chromium Controller Instrument (10x Genomics, Pleasanton, CA, USA) to generate single cell droplets containing uniquely barcoded GEMs (Gel Bead-In EMulsions). Single-cell RNA-seq libraries were obtained following the 10x Genomics recommended protocol, using the reagents included in the Chromium Single Cell 3′ v3 Reagent Kit. The libraries were sequenced on the NovaSeq S2 flow cell (Illumina, 100 cycles).

### Single-cell RNA-sequencing data analysis

Sample demultiplexing, STAR alignment to the mm10 transcriptome, filtering, and UMI counting were achieved using Cell Ranger 3.0.1 (10X Genomics)^44^. The Cell Ranger cell-calling algorithm for filtering barcodes was not employed; instead, the top 14,000 cell barcodes with the highest number of UMIs for each of the 16 samples were output for custom quality control (QC) analysis.

The custom QC analysis filtered cells to have at least 800 UMIs and 400 detected genes. For samples with more than 10,000 cell barcodes after filtering, only the top 10,000 cells with the highest numbers of UMIs were retained. These steps resulted in a final dataset of 108,397 cells. Additionally, genes that did not have any expression in at least 10 cells were removed from the analysis. Mitochondrial ratio and cell complexity were not used to filter cells due to expectations regarding the cell types present in the data.

After quality control, we used the filtered cells to perform the downstream clustering and marker identification using Seurat^19^ (v3.1.1). Normalization and variance stabilization were performed with sctransform using the 3,000 most variable genes^45^. We also regressed out cell cycle phase using the difference between the S and G2M phase scores. Using the output from sctransform, integration of samples across temperatures was performed. For clustering of the cells, Seurat’s K-nearest neighbor graph-based approach was executed using the top 50 PCs and a resolution of 1.4. Visualization of the clusters was done with the Uniform Manifold Approximation and Projection (UMAP) method^20^, and cluster quality was assessed for possible cluster artifacts. Marker identification was performed using Seurat with the Wilcoxon rank sum test, allowing for the cell type identification of the 55 clusters. Following identification and removal of doublet and erythrocyte clusters, non-immune cell clusters were subset and re-clustered again as described above, except using a clustering resolution of 1.2.

Differential gene expression analysis was performed on the non-immune cells using a pseudo-bulk approach^21^. To do so, the single cell counts were aggregated to the sample level for each cluster. Differential gene expression analysis of count data was performed using the Bioconductor R package, DESeq2 (v1.26.0). Using the likelihood ratio test (LRT), we identified all genes whose expression were significantly different between at least 2 of the conditions (temperatures). Gene Ontology (GO) term over-representation analysis was performed on the groups of significantly up- or down-regulated genes. GO analysis was performed using clusterProfiler^46^ (v3.14.3) using the clusters of genes following increasing or decreasing expression trends identified using DEGreport (v1.22.0) (Pantano L (2020). DEGreport: Report of DEG analysis. R package version 1.24.1, http://lpantano.github.io/DEGreport/).

Slingshot (v1.4.0) was used to perform branched trajectory analysis on integrated data^24^. The Seurat clusters were merged for the adipocyte progenitor cells (AP_8, AP_9, AP_25, AP_26), endothelial cells (EC_4, EC_10, EC_11, EC_15, EC_23, EC_28, EC_32, and Lymph_EC_19), vascular smooth muscle cells (VSM_0, VSM_1, VSM_2 VSM_3, VSM_5, VSM_7, VSM_20, VSM_21, VSM_22, VSM_27), and the pre-adipocyte cells (VSM_AP_6 and Adipo_16). Trajectories were calculated for the merged Seurat cell clusters, in addition to the Adipocyte (Adipo_24) cluster from the first 3 PCA components, specifying the Adipocyte (Adipo_24) cluster as the end state.

For trajectory analysis of the vascular smooth muscle cells and adipocytes, the UMAP coordinates and cell clusters for the vascular smooth muscle cells and adipocytes (VSM_0, VSM_1, VSM_2 VSM_3, VSM_5, VSM_7, VSM_20, VSM_21, VSM_22, VSM_27, VSM-AP_6, Adipo_16, and Adipo24) were used to calculate the lineages /pseudotime curves. No start or end states were specified, resulting in five separate trajectories for VSM/Adipocyte cells. VISION^25^ was used for annotating the sources of variation in the scRNA-seq data using Hallmark gene set signatures from MSigDB^47,48^.

### Flow cytometry and FACS

Adipocytes and SVF were isolated form BAT and ingWAT of Trpv1^cre^ Rosa26^mTmG^ as described above. Adipocytes were collected from the top layer, filtered through a 200 µm cell strainer, washed with 3% BSA in PBS, and centrifuged at 30 g for 5 minutes. The washing step was repeated three times and adipocytes were immediately used for FACS. The SVFs were stained with APC-conjugated Pdgfra antibody (Invitrogen). A BD FACSAria (Becton Dickinson) was used to detect endogenous membrane tagged GFP, tdTomato, and APC signals. Data were collected using DIVA (Becton Dickinson) software and analyzed using FlowJo software (Tree Star, Inc.). Debris and dead cells were excluded by forward and side scatter gating.

After sorting, adipocytes and SVFs were lysed TriPure RNA Isolation Reagent (Sigma-Aldrich) and used for RNA isolation and gene expression analysis.

### RNA isolation and quantitative RT-PCR

RNA was isolated from both adipocytes and SVF samples using phenol-chloroform extraction and isopropanol precipitation. Quantitative reverse transcription-PCR (qRT-PCR) assays were performed using an ABI Prism 7900 sequence-detection system using SYBR (Roche Applied Science, Indianapolis, IN). Relative mRNA expression was calculated by the ΔCt method and the values were normalized to the expression of ARBP. The sequences of primers are provided in Extended Data Table 1.

### Imaging of Whole Mounted Adipose Tissue

Adipose depots were dissected and cut into ∼1.5×1.5 cm pieces and were immediately mounted onto microscope slides with water. The slides were imaged on a Zeiss LSM 710 NLO confocal microscope. Images were taken with either 40X or 20X objectives. GFP and tdTomato signals were quantified using ImageJ software.

### Immunohistochemistry

BAT was fixed in 10% formalin and embedded in paraffin. 5 µm sections were prepared and stained for GFP and Plin1. Briefly, sections were deparaffinized and rehydrated, followed by an antigen retrieval step in a modified citrate buffer (Dako Target Retrieval Solution, pH 6.1, Agilent). Sections were then incubated in Sudan Black (0.3% in 70% ethanol) to reduce autofluorescence signal. Blocking was performed in Millipore blocking reagent (EMD Millipore), followed by incubating the section in Anti-GFP (ab13970, Abcam) and Anti-Perilipin-1 (D1D8 #9349, Cell Signaling Technology) overnight at 4°C. The next day, slides were washed in PBST (0.1% Tween 20 in PBS) and incubated with appropriate secondary antibodies (Invitrogen) at 1:200 dilution. The slides were mounted in mounting media with DAPI. Images were collected on a Zeiss LSM 710 NLO confocal microscope.

### Statistical analysis

Statistical methods were not used to predetermine sample size. The experiments were not randomized, and investigators were not blinded in experiments. All statistics were calculated using Graphpad Prism and RStudio. In Imaging and qPCR experiments, statistical significance was determined by Student’s t test.

### Code availability

Sample scripts used to process the data will be deposited at a publicly available Github page.

## Data Availability

The authors declare that the data supporting the findings of this study are available within the paper and its supplementary information files. scRNA-sequencing data will be deposited at the Gene Expression Omnibus (GEO).

## Acknowledgments

This work was supported in part by US National Institutes of Health (NIH) grants R01DK077097 and R01DK102898 (to Y.-H.T.), T32DK007260, F32DK102320 and K01DK111714 (to M.D.L), P30DK036836 (to Joslin Diabetes Center’s Diabetes Research Center, DRC) from the National Institute of Diabetes and Digestive and Kidney Diseases, grant #1-18-PDF-169 (to F.S.) from American Diabetes Association, and grant # 2019-02454 from CZI Seed Networks (to Y.-H.T.). Work by MP at the Harvard Chan Bioinformatics Core was funded by the Harvard Stem Cell Institute’s Center for Stem Cell Bioinformatics. We thank Alison Marotta for technical assistance.

## Author Contributions

F.S. and Y.H.T. designed and conducted the single-cell RNA-sequencing study. F.S., M.D.L., and Y.H.T. designed and conducted the lineage tracing experiments and wrote the manuscript. M.P. conducted the single-cell RNA-sequencing analyses. L.L.H. provided assistance in single-cell isolation and library preparation. T.L.H. provide technical assistance. All authors read and approved the final manuscript.

## Additional Information

Supplementary Table 1. Marker genes for clusters of cells in BAT-SVF.

Supplementary Table 2. List of differentially expressed transcripts in Pdgfra+ APCs (AP_8 and AP_9), capillary endothelial cells (EC_4), VSMs (all VSM clusters), and Schwann cells (Schwann cells_14) identified using the likelihood ratio test (LRT).

